# The furin cleavage site is required for pathogenesis, but not transmission of SARS-CoV-2

**DOI:** 10.1101/2025.03.10.642264

**Authors:** Angelica L. Morgan, Michelle N. Vu, Yiyang Zhou, Kumari G. Lokugamage, William M. Meyers, R. Elias Alvarado, Yani Ahearn, Leah K. Estes, Jessica A. Plante, Bryan A. Johnson, Mehul S. Suthar, David H Walker, Ken S. Plante, Vineet D. Menachery

**Author notes:** Equal Contribution (Angelica L. Morgan, Michelle N. Vu). Corresponding Author: Vineet D. Menachery **Email:**. ALM and MNV contributed equally to this work. Author order was determined alphabetically.

## Abstract

The SARS-CoV-2 spike, key to viral entry, has two features that differentiate it from other sarbecoviruses: the presence of a furin cleavage site (FCS; PRRAR sequence) and an extended S1/S2 loop characterized by an upstream QTQTN amino acid motif. Our prior works show that shortening the S1/S2 loop by deleting either the FCS (ΔPRRA) or deleting an upstream sequence (ΔQTQTN), ablates spike processing, alters host protease usage, and attenuates infection *in vitro* and *in vivo*. With the importance of the loop length established, here we evaluated the impact of disrupting the FCS, but preserving the S1/S2 loop length. Using reverse genetics, we generated a SARS-CoV-2 mutant that disrupts the FCS (PQQAR) but maintains its extended S1/S2 loop. The SARS-CoV-2 PQQAR mutant has reduced replication, decreased spike processing, and attenuated disease *in vivo* compared to wild-type SARS-CoV-2. These data, similar to the FCS deletion mutant, indicate that loss of the furin cleavage site attenuates SARS-CoV-2 pathogenesis. Importantly, we subsequently found that the PQQAR mutant is transmitted in the direct contact hamster model despite lacking an intact FCS. However, competition transmission showed that the mutant was attenuated compared to WT SARS-CoV-2. Together, the data argue that the FCS is required for SARS-CoV-2 pathogenesis but is not strictly required for viral transmission.

## Introduction

SARS-CoV-2 emerged in late 2019 and initiated a global pandemic with a massive impact on the economy and global public health^1^. Over the past five years, SARS-CoV-2 has continued to cause infection and disease despite development of effective vaccine and therapeutics^2^. The continued infection and spread of SARS-CoV-2 have been attributed to its ability to evolve and produce variants of concern that evade key aspects of immunity and improve viral fitness^3,4^. Much of the evolution of the virus has occurred in the spike protein with new variants able to evade host immunity^5^. In addition, several spike mutations have improved fitness and transmission of the virus in the human populations^6–9^. Despite all the changes that have occurred in SARS-CoV-2, two key elements of the spike protein have remained intact throughout the pandemic: the presence of a furin cleavage site (FCS) and an extended S1/S2 loop^10–12^. Each are unique features of SARS-CoV-2 compared to other sarbecoviruses and play key roles in infection.

The CoV spike protein consists of a globular head (S1 subunit) and a stalk (C-terminal of S1 and S2 subunit)^13^. The head is responsible for attachment and binding to host receptor ACE2. In contrast, the stalk contains the highly conserved fusion machinery. CoV spike must undergo two cleavages to enable host cell fusion, first at an S1/S2 junction site and a second at an S2’ site in the stalk to activate fusion. Unlike other sarbecoviruses, SARS-CoV-2 spike contains a unique furin cleavage site (FCS; RXXR) at the S1/S2 junction that aids in the efficiency of its cleavage. The FCS, not shared among other group 2B coronaviruses, is maintained in all SARS-CoV-2 variants. The multi-basic cleavage motif, PRRAR, exists on the S1/S2 junction loop and is involved in spike cleavage as a virion exits a producer cell. Our lab has previously investigated the importance of the FCS by generating an FCS deletion SARS-CoV-2 mutant (ΔPRRA; **Fig.1A**) and performing *in vitro* and *in vivo* characterization^11^. Our results indicated that deleting the FCS attenuates SARS-CoV-2 infection in respiratory cells and pathogenesis *in vivo* compared to wild-type (WA-1 strain)^11^.

**Figure 1.**
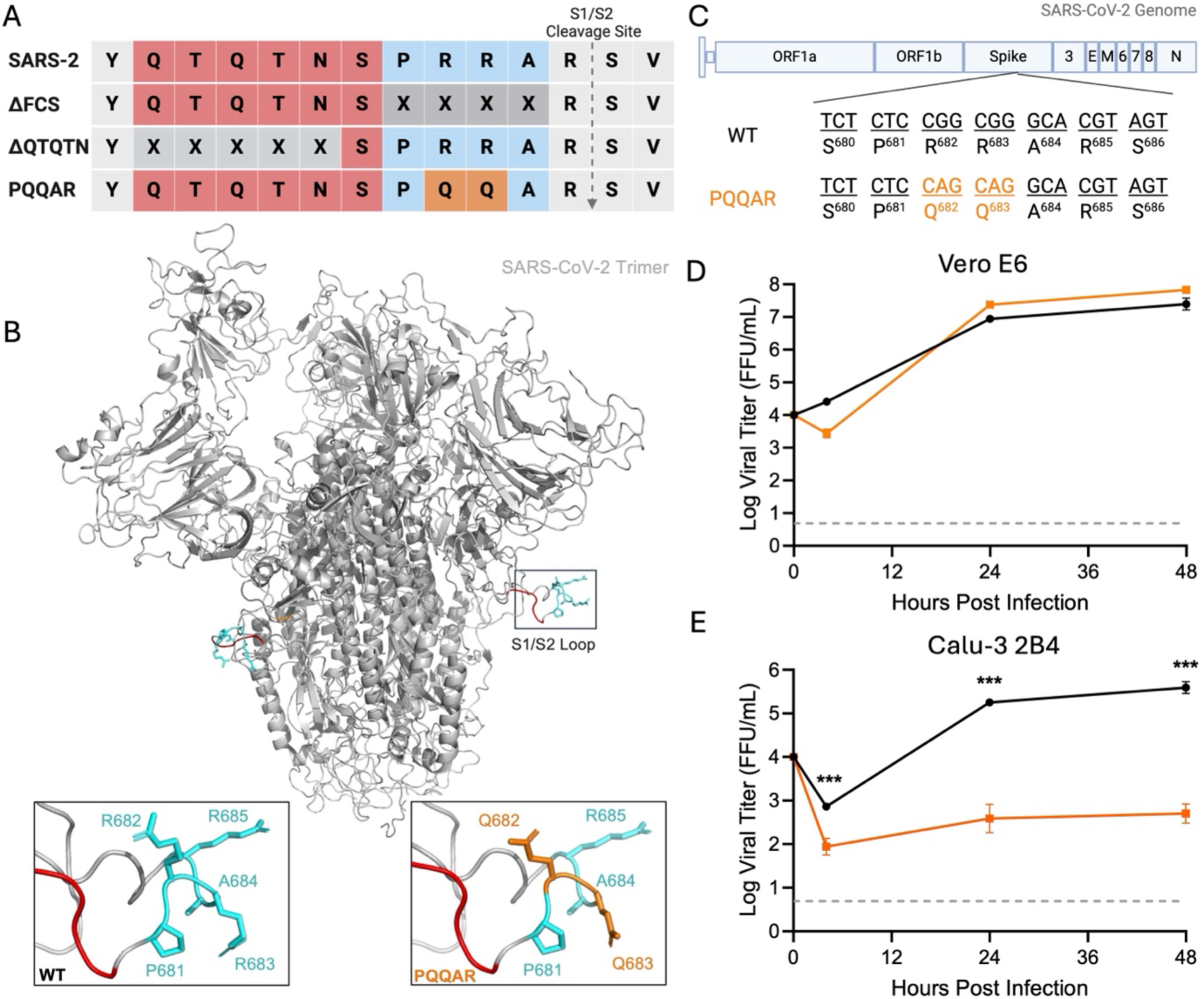
Generation and *in vitro* characterization of SARS-CoV-2 PQQAR mutant. (A) Alignment of the S1/S2 cleavage site of SARS-CoV-2 WA1 and series of mutant viruses generated for evaluation including deletion of the furin cleavage site (ΔFCS), truncation of the extended loop (ΔQTQTN), and disruption of the furin cleavage site motif (PQQAR). (B) SARS-CoV-2 spike trimer structure (gray) highlighting the S1/S2 cleavage loop. WT (left) and PQQAR mutant (right) are zoomed with mutated residues (Q682, Q683) in orange to disrupt the furin cleavage site. (C) Schematic of SARS-CoV-2 spike with PQQAR substitutions identified. (D) Viral titer from Vero E6 infected with WT (black) or PQQAR (orange) SARS-CoV-2 at an MOI of 0.01 (n = 3). (E) Viral titer from Calu-3 2B4 infected with WT or PQQAR SARS-CoV-2 at an MOI of 0.01 (n = 3). Data are mean ± SD. Statistical analysis measured by two-tailed Student’s t test. *P ≤ 0.05; **P ≤ 0.01; ***P ≤ 0.001

Soon after, we investigated the function of the QTQTN motif immediately upstream the FCS, which is commonly deleted in cell culture stocks. Our group generated a QTQTN deletion mutant (ΔQTQTN; **Fig. 1A**) and performed *in vitro* and *in vivo* testing^12^. The ΔQTQTN mutant, like ΔPRRA, has attenuated infection in respiratory cells and pathogenesis *in vivo*^12^. Both deletion mutants shorten the spike S1/S2 loop and demonstrate that loop length is important for SARS-CoV-2 pathogenesis. However, these studies do not evaluate importance of the FCS in the context of the extended S1/S2 loop found in SARS-CoV-2. To address this question, we generated an infectious clone of SARS-CoV-2 that disrupts the FCS without shortening the loop (PQQAR mutant). We demonstrate that disruption of the FCS attenuates SARS-CoV-2 replication in human respiratory cells and pathogenesis in hamsters. The PQQAR mutant has inhibited spike processing and altered protease usage for host cell entry. We also found that an intact FCS is not required for SARS-CoV-2 contact transmission in hamsters, but plays a role in transmission efficiency. Together, the data indicate that the SARS-CoV-2 FCS is important for SARS-CoV-2 infection, pathogenesis, and transmission.

## Results

### Generation of the PQQAR mutant

The furin cleavage site (FCS; PRRA motif) of SARS-CoV-2 exists at the S1/S2 junction site on an external disordered loop in the spike protein (**Fig. 1B**). Our prior work evaluated the importance of the furin cleavage site by deleting the PRRA motif upstream of the S1/S2 cleavage site^11^. While the ΔPRRA mutant was attenuated, subsequent work shortening the S1/S2 loop (ΔQTQTN) of SARS-CoV-2 had a similar phenotype, indicating the importance of the loop length in SARS-CoV-2 pathogenesis^12^. However, the role of the actual FCS in the context of the extended S1/S2 spike loop was still unclear. To investigate how the FCS, independent of loop length, impacts infection and pathogenesis, we generated an FCS mutant in the SARS-CoV WA-1 backbone that substituted arginine at position 682 and 683 to glutamines (PQQAR) (**Fig. 1C**). The PQQAR mutant SARS-CoV-2 maintains the S1/S2 loop length but disrupts the furin cleavage site allowing evaluation of the role of the FCS. Using our reverse genetic system, we were able to recover the PQQAR mutant which grew to robust stock titers in Vero cells.

### PQQAR mutant attenuates viral replication in respiratory cell

Deletion of the FCS and shortening of the S1/S2 loops of the spike had previously been shown to impact viral replication^11,12^. Therefore, we first examined the PQQAR mutant in Vero E6 cells (African green monkey kidney cells) which lack type I interferon responses^14^. Following low MOI (0.01) infection, the PQQAR mutant shows no significant attenuation compared to WT SARS-CoV-2 in Vero cells (**Fig. 1D**). In fact, the PQQAR mutant grew to slightly higher viral titer, although not significantly different that WT. These results are consistent with prior studies shortening the SARS-CoV-2 S1/S2 loop length and correspond with a fitness advantage in Vero cells^11,12^. In contrast, the PQQAR mutant has attenuated replication in Calu-3 2B4 cells, a human respiratory cell line (**Fig. 1E**). At 24 and 48 hours post infection (hpi), the PQQAR mutant has ∼3 log reduction in titers compared to WT. Again, these results are consistent with prior studies(ΔQTQTN, ΔPRRA)^11,12^ and demonstrate that the both the S1/S2 loop and the FCS play a role in effective SARS-CoV-2 replication in human respiratory cells.

### PQQAR mutant has attenuated *in vivo* pathogenesis

Having established attenuation in human respiratory cells, we next evaluated how disruption of the FCS impacts SARS-CoV-2 pathogenesis *in vivo*. Utilizing the golden Syrian hamster model, three- to four-week-old male hamsters were intranasally infected with 10^5^ focus forming units (FFU) of WT or PQQAR mutant SARS-CoV-2 and followed over a 7-day time course (**Fig. 2A**). WT-infected hamsters exhibited weight loss starting at 2 dpi with peak weight loss of ∼10-12% before beginning to recover at 6 dpi (**Fig. 2B**). In contrast, the PQQAR mutant produced minimal weight loss following infection. Similarly, disease score corresponded with weight loss as WT had observable disease between days 3 and 6 characterized by ruffled fur, hunched posture, and reduced activity. In contrast, no disease was noted in PQQAR mutant infected hamsters. Together, the data demonstrated that disruption of the FCS attenuated disease in the hamster model of SARS-CoV-2 infection.

**Figure 2.**
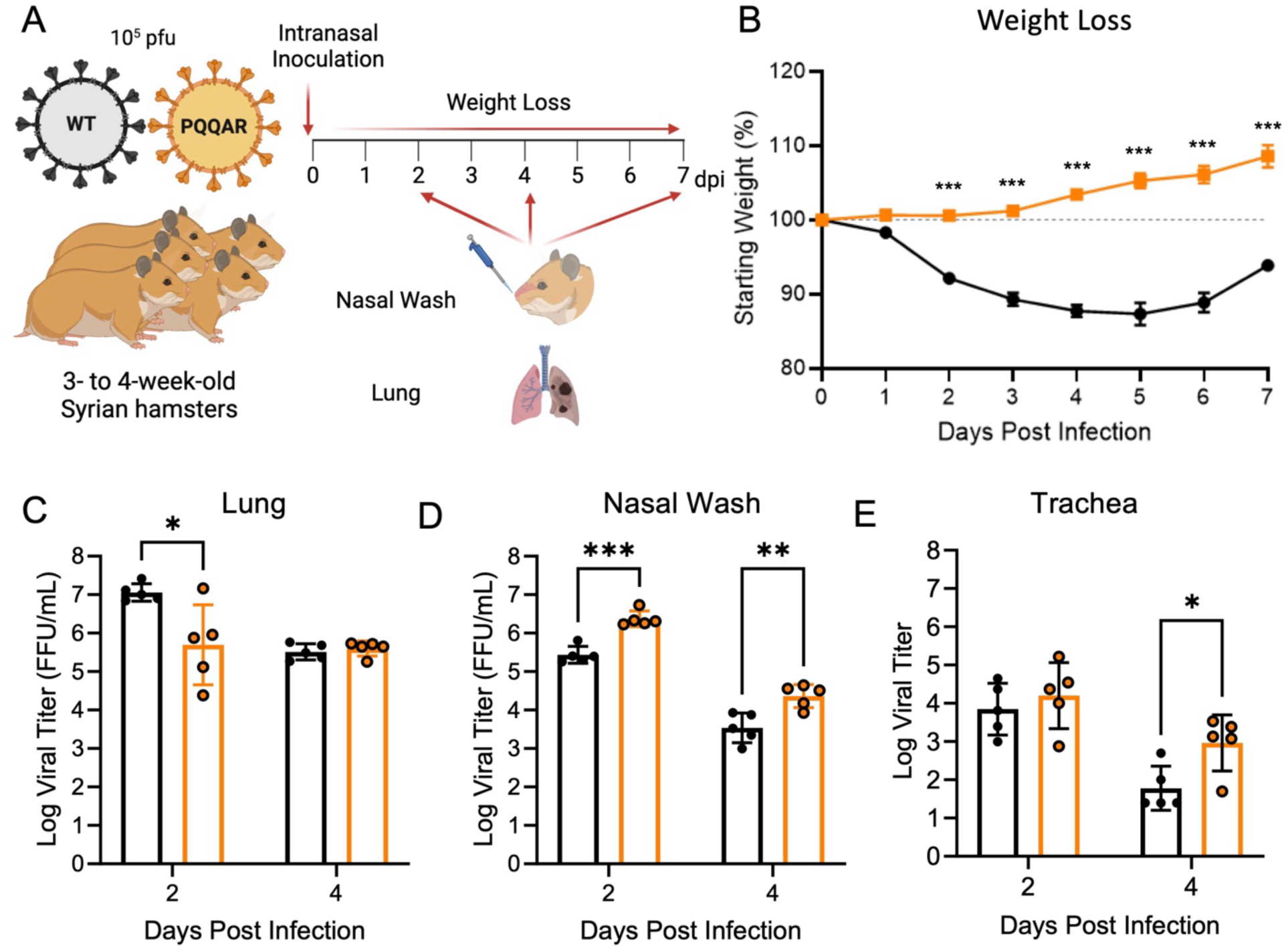
SARS-CoV-2 PQQAR mutant attenuated in golden Syrian hamsters. (A-B) Schematic of golden Syrian hamster infection with WT (black) or PQQAR mutant (orange) SARS-CoV-2. Three- to four-week-old male hamsters were infected with 10^5^ pfu and monitored (B) weight loss and disease for 7 days post infection. (C-E) Viral titers were measured at days 2 and 4 from (C) infected lung, (D) nasal wash, and (E). Data are representative of mean ± SEM. Statistical analysis measured by two-tailed Student’s t test. *P ≤ 0.05; **P ≤ 0.01; ***P ≤ 0.001. Experimental schematic made in Biorender.

### PQQAR mutant has altered replication in upper and lower respiratory tract

Having established attenuated disease, we next evaluated viral replication in the lung, nasal wash, and trachea following infection with WT and PQQAR mutant. Examining the lung, we found that both WT and the PQQAR mutant replicated in the lungs following infection (**Fig. 2C**). However, the PQQAR mutant had a 1 log reduction in viral replication compared to WT at day 2 but was equal to WT at day 4. Probing the upper airway, the PQQAR mutant had a significant increase (∼10-fold) in viral replication from nasal washes at both days 2 and 4 as compared to WT (**Fig. 2D**). These results are similar to prior studies that shortened the spike S1/S2 loop length or deleted the FCS^11,12^. In addition, the PQQAR mutant had similar and increased replication in the trachea at days 2 and 4 as compared to WT-infected hamsters (**Fig. 2E**). Overall, the replication data demonstrate viral replication attenuation of the PQQAR mutant in the lung but also augmented replication of the mutant in the upper airway tissues.

### Viral antigen staining confirms attenuation of PQQAR mutant *in vivo*

To further evaluate viral replication and distribution, we examined antigen staining following infection with WT or PQQAR mutant at days 2 and 4 post infection. Briefly, lung sections from infected hamsters were stained for SARS-CoV-2 nucleocapsid and scored in a blinded manner for antigen in the airways, parenchyma, and overall as previously described^8^. We observed wide-spread antigen staining in both the parenchyma and airways following WT SARS-CoV-2 infection at both day 2 and 4 (**Fig. 3A-B**). In contrast, the PQQAR mutant had more limited antigen staining compared to WT (**Fig. 3C-D**). While showing modest attenuation in the airways (**Fig. 3E**), the scores from the parenchyma (**Fig. 3F**) and overall (**Fig. 3G**) demonstrated that the PQQAR mutant had a significant deficit in antigen staining as compared to WT SARS-CoV-2 infection. The antigen staining in the large airways may be consistent with potentially greater replication in the airways (**Fig. 2D-E**). Similarly, the attenuation in the parenchyma and overall staining corresponds to reduced viral titer in the lung of PQQAR mutant-infected animals (**Fig. 2C**). Together, these results demonstrate attenuation in PQQAR mutant as compared to WT in terms of viral infection and distribution in the lung.

**Figure 3.**
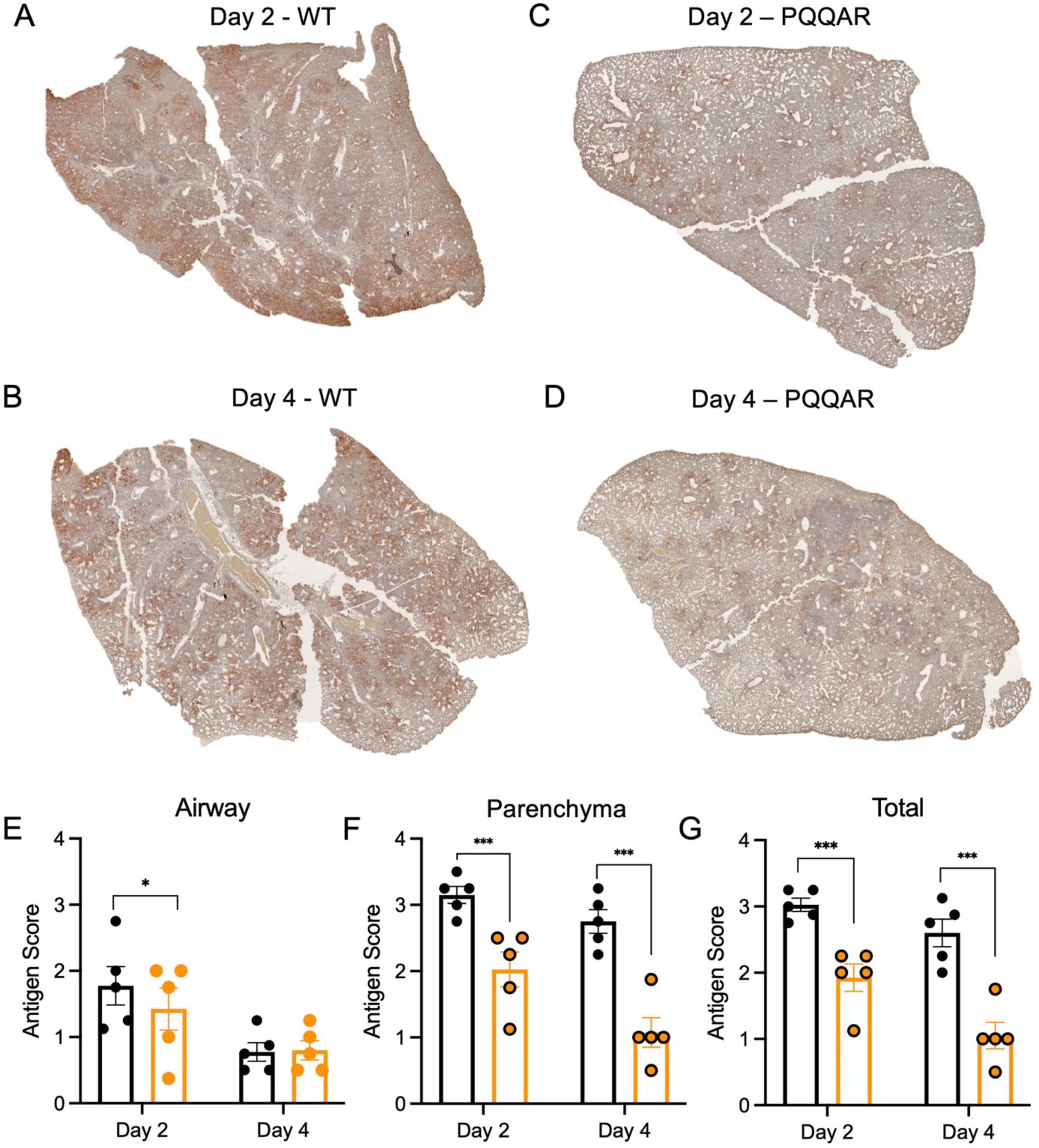
Attenuated antigen staining in PQQAR-infected hamsters. (A-D) Nucleocapsid antigen staining of left lung section from hamsters infected with 10^5^ ffu of either (A-B) WT or (C-D) PQQAR mutant at 2 or 4 dpi. Antigen staining was scored in a blinded manner by location in the (E) airway, (F) parenchyma, and (G) total for WT (black) or PQQAR (orange) infected lungs. Statistical analysis measured by two-tailed Student’s t test. *P ≤ 0.05; **P ≤ 0.01; ***P ≤ 0.001

### PQQAR mutant infection shows reduced immune infiltration and damage

To further evaluate disease and damage, WT- and PQQAR-infected hamsters were evaluated for changes in histopathology. Lung sections were H&E stained and subsequently evaluated by a board-certified pathologist in a blinded manner. After infection, WT-infected hamsters demonstrated significant disease and damage characterized by interstitial pneumonia, peribrochiolitis, epithelial cytopathology, arterial mononuclear cell margination, and presence of polymorphonuclear cells in the bronchioles in some animals at day 2 (**Fig. 4A**). In contrast, significantly less disease was observed in the PQQAR mutant infected hamsters and was characterized by mild bronchiolitis, and interstitial pneumonia (**Fig. 4B**). At day 4, WT infected hamsters showed increased lung involvement by area involved with the addition of hemorrhage, pulmonary edema, perivasculitis, and infiltration of mononuclear cells (**Fig. 4C**). Notably, while increased compared to day 2, PQQAR mutant-infected animals had much less lung involvement and less severe disease and damage as compared to WT (**Fig. 4D**). While no significant disease and damage was observed in mock infected animals (**Fig. 4E),** the SARS-CoV-2 associated lesion were much more extensive in WT as compared to PQQAR mutant infected hamsters. WT-infected hamsters had significant increases in SARS-CoV-2 lesion percentages in lung section at days 2, 4, and 7 days post infection as compared to PQQAR mutant infection (**Fig. 4F**). These results are consistent with the weight loss and disease data (**Fig. 2B**) and indicate that the PQQAR mutant is attenuated in terms of pathogenesis *in vivo*.

**Figure 4.**
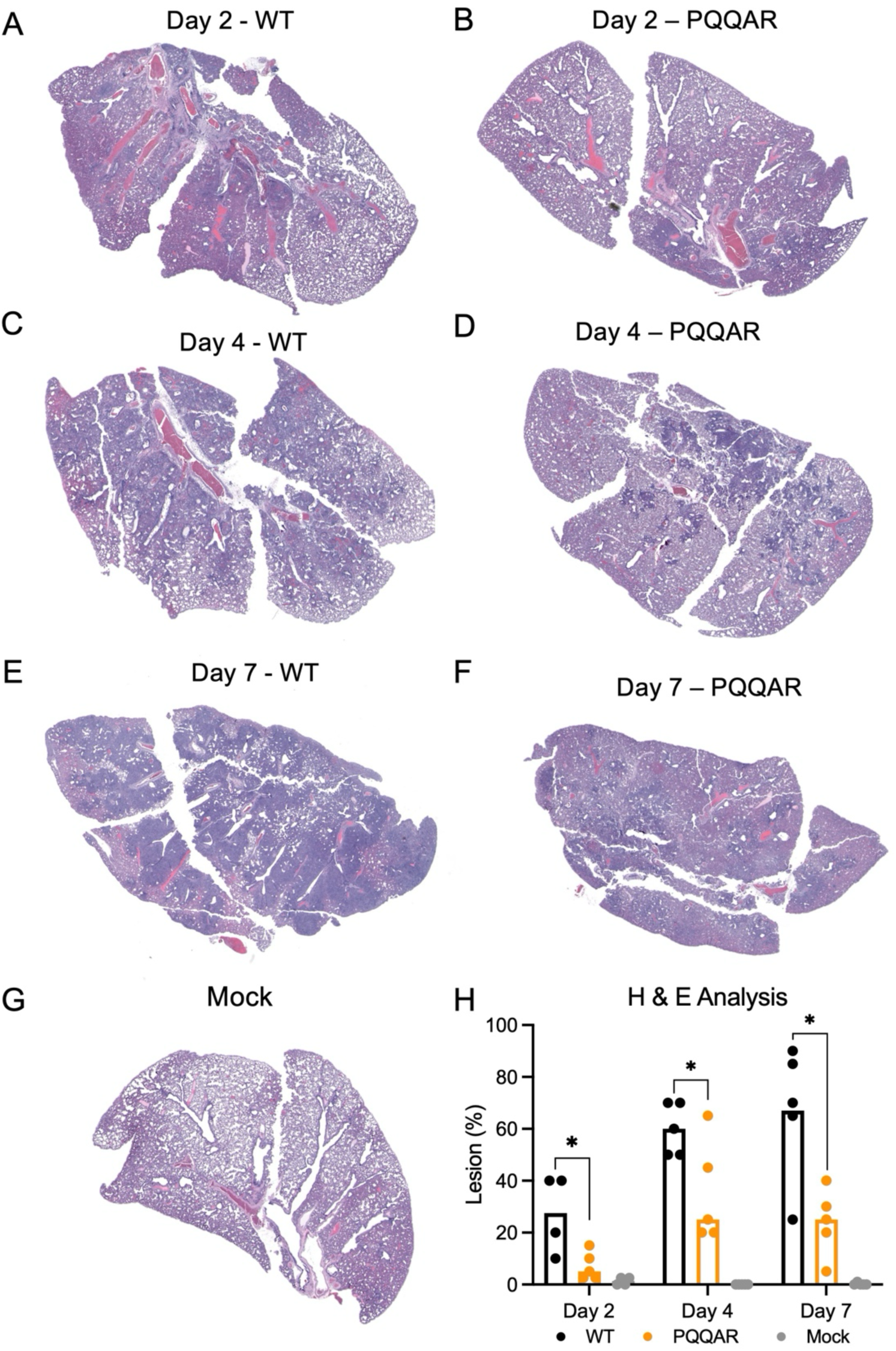
Reduced inflammation and damage in PQQAR-infected lungs. (A-G) Representative H&E staining of left lung of hamsters infected with 10^5^ pfu of either WT or PQQAR SARS-CoV-2 at (A-B) 2 days, (C-D) 4 days, or (E,F) days post infection or (G) mock. (H) WT (black), PQQAR R(orange), or PBS (grey) lung sections from each day were scored for histopathological analysis with sections from individual animal averaged and representing a single point. Statistical analysis measured by two-tailed Student’s t test. *P ≤ 0.05; **P ≤ 0.01; ***P ≤ 0.001

### FCS disruption attenuates spike processing

Spike mediated entry requires sequential cleavage at the S1/S2 junction and the S2’ site to activate fusion^15^. Prior work has established that the SARS-CoV-2 spike is partially cleaved prior to release from cells^16^; shortening the S1/S2 loop disrupts this cleavage on released virions. Here, we evaluate if FCS disruption affected spike cleavage on purified virions from the PQQAR mutant. Briefly, WT, PQQAR, and the ΔFCS mutant SARS-CoV-2 were collected from Vero E6 cells, purified using ultracentrifugation sucrose cushion, inactivated, and blotted for spike and nucleocapsid (**Fig. 5A**). Normalizing to nucleocapsid, we observe that both PQQAR and ΔFCS mutants have a reduction in S1/S2 cleavage product as compared to WT (**Fig. 5B**). While WT SARS-CoV-2 has ∼40% of spike cleaved following infection of Vero cells, both disruption and deletion of the FCS results in mutants with very minimal S1/S2 cleavage product (**Fig. 5C**). Together, the results indicate that an intact FCS is required for spike processing prior to release of SARS-CoV-2 virions.

**Figure 5.**
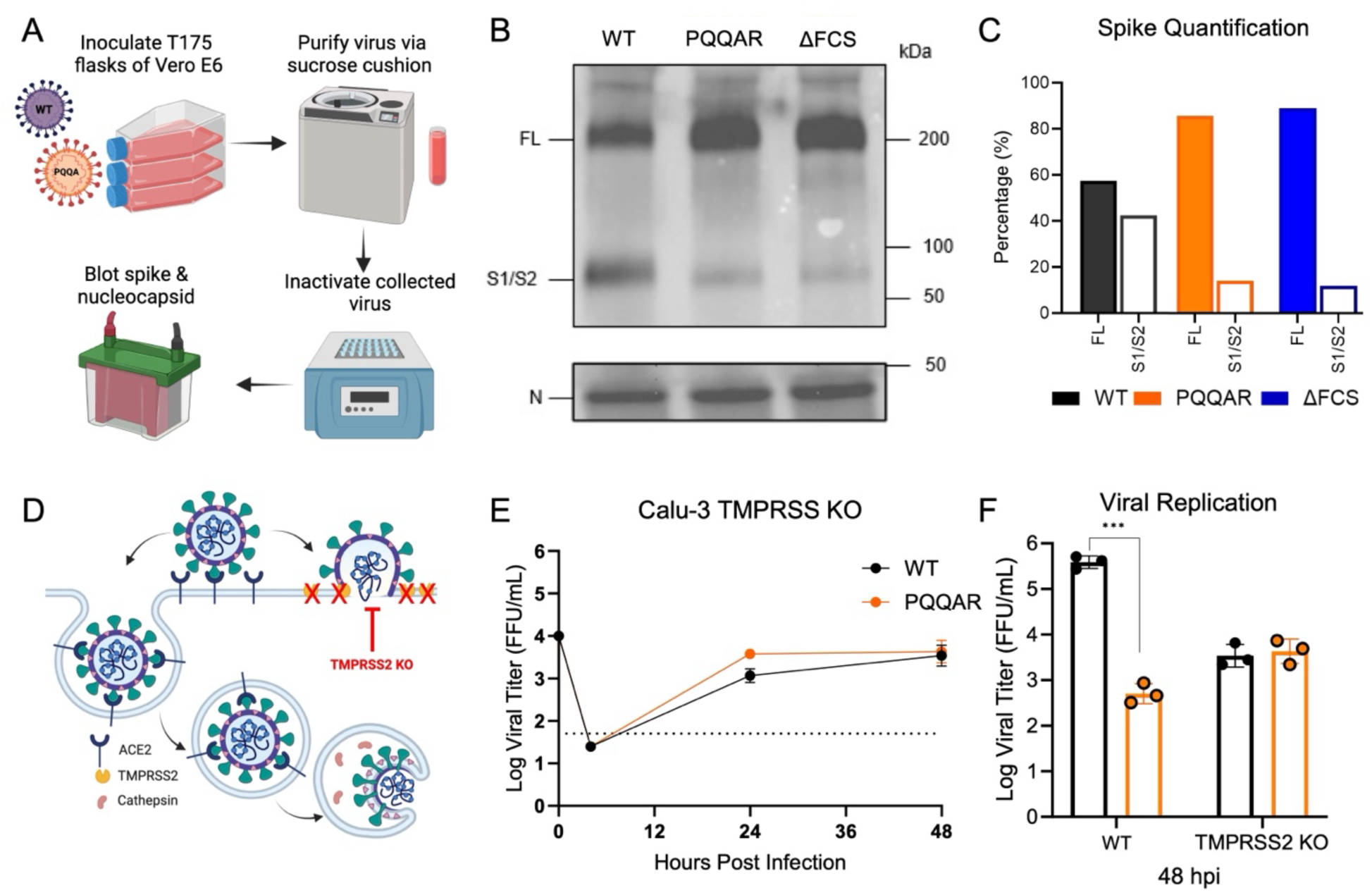
Disruption of FCS alters spike processing and protease usage. (A) Schematic of SARS-Cov-2 virion sucrose cushion purification approach. (B) Lysates from sucrose cushion purified WT, PQQAR, and ΔFCS virions grown in Vero E6 were probed with α-Spike and α-Nucleocapsid (N) antibodies by Western blot. Full-length spike (FL) and S1/S2 cleavage product are indicated. (C) Quantification of densitometry of the proportion between FL (black) and S1/S2 (red) of the total spike shown (lower). (D) Schematic of SARS-CoV-2 entry and protease usage including knockout of TMPRSS2 mediated entry. (E) Viral titer from Calu-3 TMPRSS2 knock-out cells infected with WT (black) or PQQAR (orange) SARS-CoV-2 at an MOI of 0.01 (n = 3). (F) Viral titer at 48hpi from Calu3 WT (Fig. 1D) and Calu3 TMPRSS2-/- cells. Statistical analysis measured by two-tailed Student’s t test. *P ≤ 0.05; **P ≤ 0.01; ***P ≤ 0.001. Entry schematic made in Biorender.

### TMPRSS2 plays role in attenuation of PQQAR mutant

SARS-CoV-2 enters the host cell through either the endosomal route mediated by host cathepsins or the cell surface route mediated by host serine proteases like TMPRSS2 (**Fig. 5D**)^15^. The PQQAR mutant shows significant attenuation in Calu-3 2B4, but has no deficit in Vero E6 cells (**Fig. 1C, 1D**). A major distinction between the two cell lines is expression of TMPRSS2, which is highly expressed in Calu3 cells, but absent in Vero E6 cells. To determine if TMPRSS2 expression contributes to attenuation, we utilize Calu3 cells with TMPRSS2 knocked out. We subsequently infected the TMPRSS2 KO Calu3 cells with both WT SARS-CoV-2 and PQQAR mutant as previously described. Following infection, we note that both PQQAR and WT virus replication is not significantly different over the course of infection (**Fig. 5E**). Compared to standard Calu3 cells (**Fig. 5F**), WT SARS-CoV-2 infection was reduced to the same level of PQQAR mutant indicating that the primary difference between replication of the WT and FCS mutant virus in Calu3 cells is due to the activity of TMPRSS2. These results indicate that spike processing through the FCS plays an important role in utilization of TMPRSS2 mediated entry of SARS-CoV-2.

### Intact FCS is not required for transmission of SARS-CoV-2 *in vivo*

The disruption of the FCS by the PQQAR mutant demonstrates that the importance of the motif to SARS-CoV-2 infection of respiratory cells and pathogenesis *in vivo*. Yet, it remains unclear if the furin cleavage site is required for transmission of SARS-CoV-2. While attenuated in the lung, the PQQAR mutant displayed robust and augmented replication in the upper airways as compared to WT SARS-CoV-2 (**Fig. 2C**). With the upper airway infection thought to seed transmission, it left the possibility that the PQQAR mutant could still be transmited without a functional FCS. To test this question, we performed studies with WT and PQQAR mutant SARS-CoV-2 to evaluate the capacity of the mutant virus to be transmitted in hamsters. Briefly, “donor” hamster were intranasally infected with 10^5^ (FFU) of WT SARS-CoV-2 or PQQAR mutant (**Fig. 6A**). After 24 hours, each donor was individually co-housed 1:1 with a naïve “recipient” with no additional barriers to evaluate contact transmission for 8 hours. After cohousing, donors and recipients were separated into individual cages for the duration of the study. Before being returned to new cages, donor hamsters were nasal washed to measure their shedding titer on the day of exposure. Animals were subsequently examined for changes in weight and disease, euthanized two days post infection/exposure, and tissues/washes evaluated for viral load.

**Figure 6.**
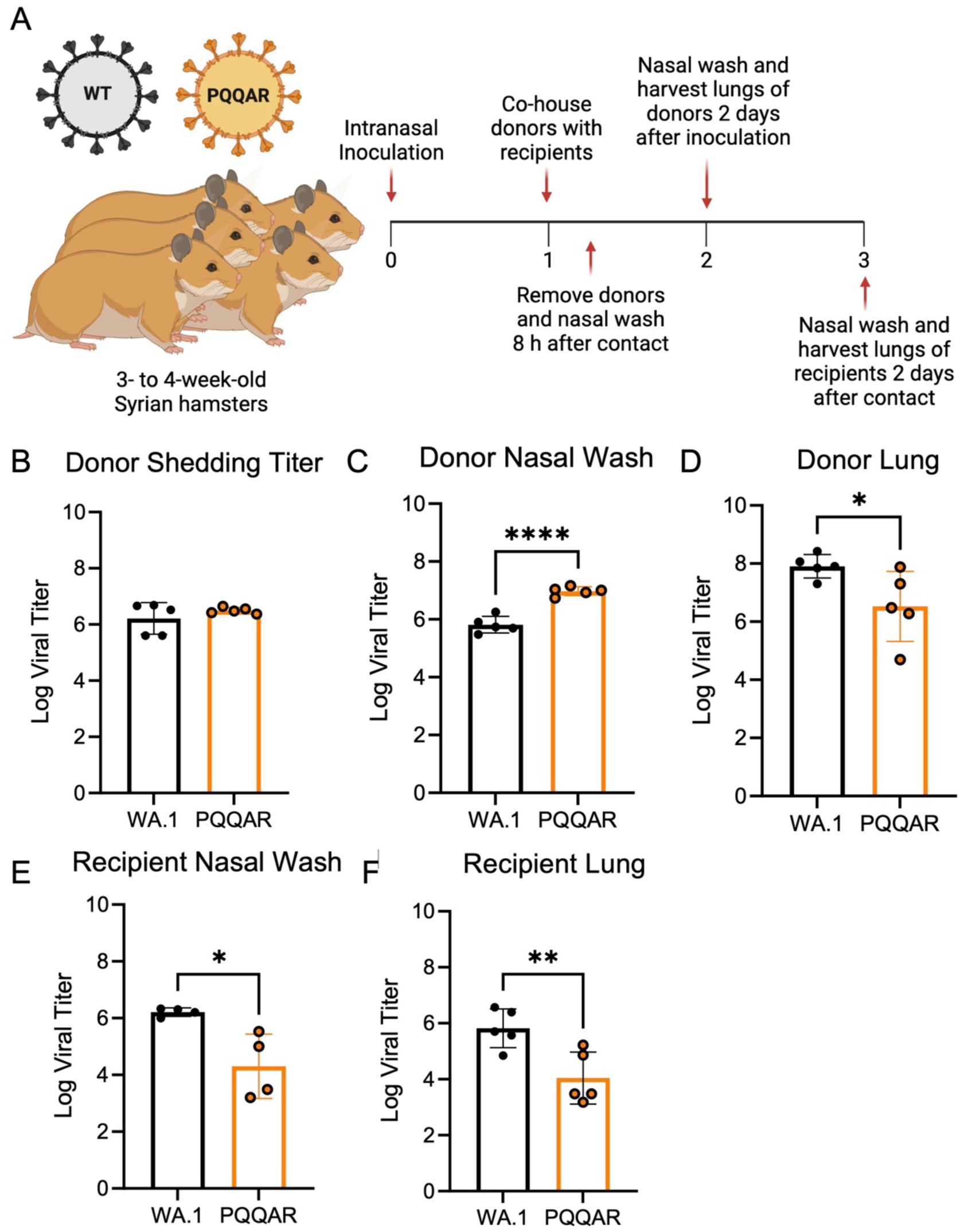
The FCS is not required for SARS-CoV-2 transmission. (A) Schematic of transmission experiment in golden Syrian hamsters. Three- to four-week-old male donor hamsters were intranasally infected with 10^5^ pfu of WT or PQQAR SARS-CoV-2 and individually housed. Donors were subsequently paired 1:1 with recipients 24 hpi and cohoused for 8 hours before separating and nasal washing donors. (B-F)Nasal washes and lungs were collected at 2 days post infection for donors (dpi) (B-D) and post contact for recipients (E-F). Viral titers were measured using focus forming assays for donor and recipient samples. Statistical analysis measured by two-tailed Student’s t test. *P ≤ 0.05; **P ≤ 0.01; ***P ≤ 0.001. Experimental schematic made in Biorender.

Following infection of donors, we observed that the shedding titers in the nasal wash were similar between WT and PQAAR mutant after the exposure period (**Fig. 6B**). We also observed that donor titers at 2 days post infection showed the PQAAR mutant had higher viral load in the nasal wash, but reduced titer in the lung (**Fig. 6C-D**). These results are consistent with observations from acute infection showing greater replication of PQQAR mutant in the airways versus the lung (**Fig. 2C-D**). Examining transmission from donors to recipient, we observed that transmission of both WT and PQQAR mutants was 100% successful. All 5 donor pairs infected with either WT or PQQAR mutant resulted in detectable viral loads in the nasal wash and lungs of recipient animals 2 days post exposure (**Fig. 6E-F**). Yet, while transmission occurred, PQQAR mutant recipient hamsters had significant lower viral titers compared to WT for both the lung and nasal washes (**Fig. 6E-F**). The results suggest that while the FCS is not strictly required for transmission, its presence impacts the viral load observed in recipient animals. Regardless of reduced viral load, the results indicate that an intact FCS is not required for SARS-CoV-2 transmission in the direct contact hamster model.

### FCS necessary for transmission fitness *in vivo*

To investigate how loss of the FCS impacts transmission fitness, we performed a competition transmission study between WT and PQQAR mutant virus in hamsters. Briefly, donor hamsters were intranasally infected with an equal mixture (1:1) of WT and PQQAR mutant virus with a total of 10^5^ focus forming units (**Fig. 7A**). At 24 hours post infection, donor hamsters were cohoused 1:1 with naïve recipients for 12 hours before separation into individual cages. Upon separation, the donor hamsters were nasal washed after exposure. Animals were subsequently examined for weight loss and disease over a four-day time course. Animals were euthanized and RNA from lung tissue/nasal washes were collected from the hamsters at 2- and 4-days post-infection or post-contact. The extended time point (4 dpi) evaluated if the competition results changed after an prolonged period of time post infection.

**Figure 7.**
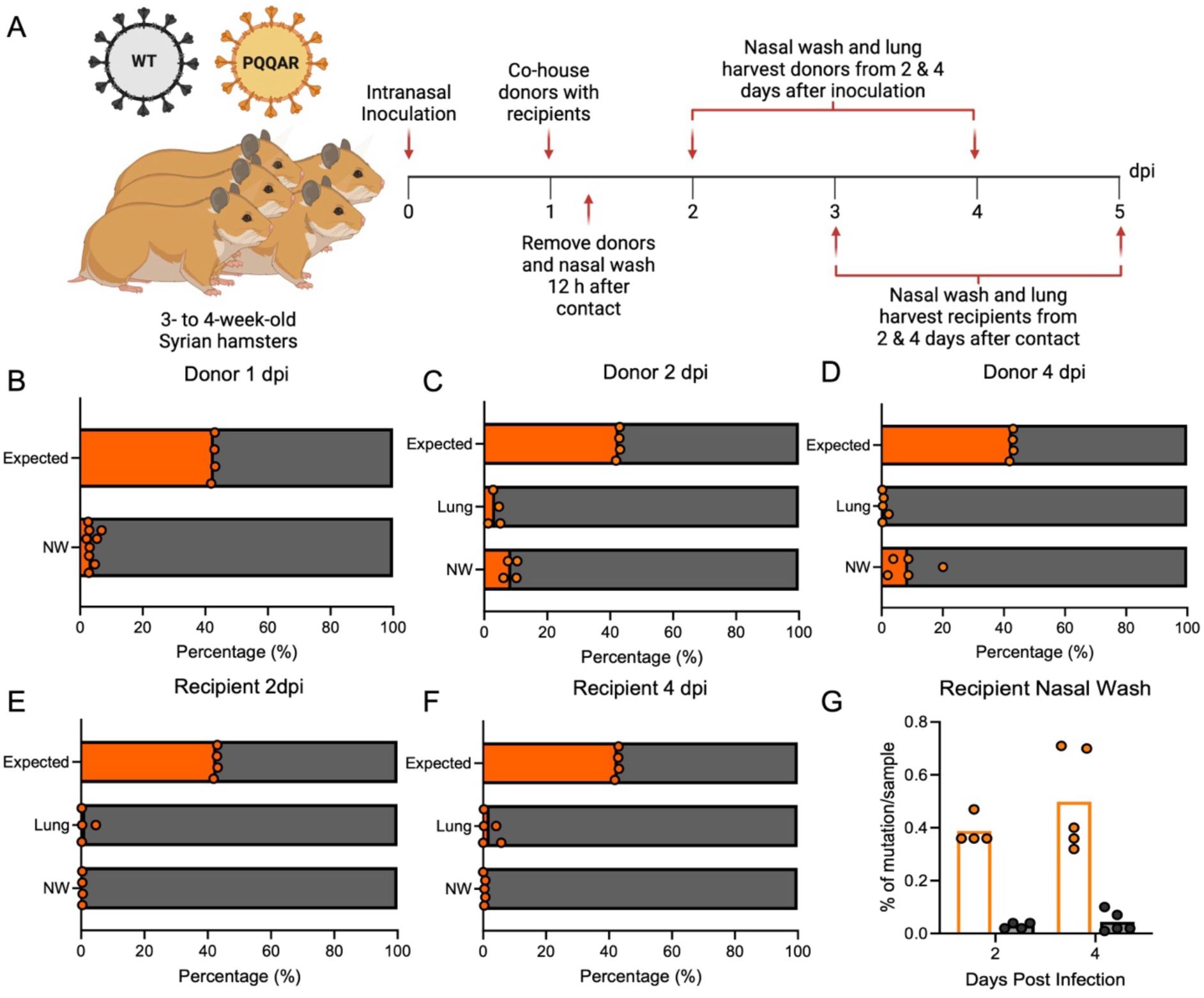
The furin cleavage site impacts SARS-CoV-2 transmission efficiency. (A) Schematic of transmission competition experiment in golden Syrian hamsters. Three- to four-week-old male donor hamsters were intranasally infected with 10^5^ pfu of WT:PQQAR SARS-CoV-2 in a 1:1 ratio and were individually housed. (B-F) After 24 hpi, donors were paired with recipients and cohoused for 12 hours before separating and nasal washing donors. Nasal washes and lungs were collected at 2 and 4 days post infection for donors (dpi) and post contact for recipients (dpc). Next generation sequencing was performed on extracted RNA to measure the percentage of WT (grey) and PQQAR (orange) present in nasal wash and lung of donors (B-D) and recipients (E-F). The expected distribution (B-F, top bar) based on NGS percentage mutant/WT observed in the inoculating dose (two inoculum preparations with RNA sequenced twice from each). Statistical analysis measured by two-tailed Student’s t test. *P ≤ 0.05; **P ≤ 0.01; ***P ≤ 0.001. Experimental schematic made in Biorender.

We used Next-Generation sequencing (NGS) to evaluate transmission dynamics of WT and PQQAR from *in vivo* competition. NGS libraries were made from extracted RNA samples with “Tiled-ClickSeq”^17^, an approach that utilizes “click chemistry” and >300 SARS-CoV-2 specific primers to fully examine the SARS-CoV-2 genome. The abundance of viral RNA comprised of WT or PQQAR genotypes suggests their relative ratio during transmission competition. Examining donor hamsters, we find that PQQAR mutant had substantially reduced transmission efficiency than WT for 1, 2, or 4 dpi. (**Fig. 7B-D**). From nasal wash, PQQAR mutant comprised ∼2-20.1% of total viral reads (mean: 8.63% +/- 0.05%) or a ∼12.3-fold transmission reduction compared to WT. From lung tissue, PQQAR mutant comprised ∼0.2-5.1% of total viral reads (mean: 1.95% +/- 0.02%) or 162.7-fold transmission reduction relative to WT. Our competition results differ from the findings in the first transmission study that found robust replication of the PQQAR mutant in donor animals (**Fig. 6B-D**). The single viral inoculum experiment (**Fig. 6**) demonstrates that PQQAR is replication-competent *in vivo*, while the mixed inoculum experiment (**Fig. 7**) shows that although PQQAR can replicate, it is outcompeted by WT SARS-CoV-2.

From recipient hamsters, the PQQAR mutant was found to have even lower transmission efficiency as compared to WT (**Fig. 7E-F**). Examination of both the lung and nasal washes revealed that the PQQAR mutant was found to be <5% of the reads. Notably, despite the low levels of viral reads, the PQQAR mutant RNA was detected in all recipient animals at levels well above the number of background mutations (**Fig. 7G**) indicating that the detected PQQAR genotype is a result of actual transmission, rather than random mutations. However, the PQQAR mutant is unable to compete with WT virus that maintains an intact FCS. Together, the results demonstrate that while not strictly required, the SARS-CoV-2 spike FCS site aids in transmission efficiency.

## Discussion

The presence of the furin cleavage site in SARS-CoV-2 spike protein plays a key role in infection and pathogenesis. As novel variants continue to emerge, SARS-CoV-2 has maintained its furin cleavage site and extended S1/S2 loop length in the spike protein indicating its necessity. While we previously established the importance of the SARS-CoV-2 S1/S2 loop length^11,12^, this study addresses how the furin cleavage site itself impacts SARS-CoV-2 infection and pathogenesis. We found that disruption of the FCS significantly attenuates replication in human respiratory cells *in vitro* and pathogenesis *in vivo*. The FCS mutant (PQQAR) has inhibited spike processing and modified host protease usage during cell entry. Notably, despite attenuated pathogenesis, the PQQAR mutant is successfully transmitted to all contact recipients but has decreased transmission fitness. Together, our data indicate that the furin cleavage site is critical for SARS-CoV-2 infection, pathogenesis, and transmission fitness.

Overall, our studies confirm that both the loop length and an intact furin cleavage site are required for pathogenesis of SARS-CoV-2. Previously, deletion mutants, ΔPRRA and ΔQTQTN, both had attenuated pathogeneicity in hamsters; the results showed that the SARS-CoV-2’s extended S1/S2 loop played a critical role in pathogenesis^11,12^. However, the previous findings failed to evaluate the FCS requirement in the context of an extended S1/S2 loop. In this study, we demonstrate that disruption of the FCS also causes attenuation of *in vivo* pathogenesis and has viral replication kinetics similar to the ΔPRRA and ΔQTQTN deletion mutants. The PQQAR mutant maintains the extended loop length, but still results in reduced body weight loss and significantly less lung pathology in hamsters. While there is slightly augmented replication in the upper airways, the PQQAR mutant infected animals had lower viral loads in the lung and reduced antigen staining. Together, these data highlight the importance of both an intact FCS and extended S1/S2 loop in SARS-CoV-2 pathogenesis.

Combined with prior studies, our results indicate that spike processing and protease usage play a key role in SARS-CoV-2 pathogenesis. Disruption of the furin cleavage site (PQQAR) or shortening of the S1/S2 loop (ΔQTQTN) both independently ablate spike processing of SARS-CoV-2 virions^11,12^. Spike processing changes on the SARS-CoV-2 virion also correspond with changes in protease usage; the FCS and shortened loop mutants were unable to utilize TMPRSS2-mediated pathways as effectively. The changes in spike processing and protease usage also correspond with attenuation of *in vivo* pathogenesis. The results argue that the S1/S2 loop length and the FCS both play critical roles in SARS-CoV-2 infection and disease. Importantly, a number of other SARS-CoV-2 spike mutations found in variants of concern have been shown to modulate both spike processing and protease usage^8,18,19^. In each case, these results impact disease pathogenesis and highlight the importance of both the FCS and S1/S2 loop in SARS-CoV-2 disease and damage.

In contrast to its role in pathogenesis, our results indicate that an intact FCS is not strictly required for SARS-CoV-2 transmission but likely plays a role in transmission and infection efficiency. Prior studies in ferrets found that deletion of the furin cleavage site ablated transmission of the virus via direct contact^20^. In contrast, we found that disrupting the FCS did not prevent transmission of SARS-CoV-2 in the direct contact hamster model with all 5 recipients infected. A major distinction is the nature of the FCS mutations as the prior study deleted the entire cleavage loop motif (PRRAR) ablating the S1/S2 site as well as shortening the loop^20^. Our study maintains the S1/S2 loop length by disrupting only the furin targeted motif and also maintaining the Sarbecovirus cleavage site (PQQA<u>R</u>)^7^. Our findings indicate that the intact FCS is not a requirement but leaves the possibility that the S1/S2 extended loop may be necessary for efficient transmission. Further studies are necessary to explore the role of the extended S1/S2 loop for SARS-CoV-2 transmission.

While the PQQAR mutant was able to be passed to 100% of recipient animals, the efficiency of transmission was reduced. Examining viral loads, the PQQAR mutant recipient had lower viral loads as compared to WT controls following single virus transmission study. Similarly, the PQQAR mutant was outcompeted by WT SARS-CoV-2 in direct competition studies (**Fig. 7**). While the results corresponded with attenuation PQQAR mutant loads in the lung upon direct challenge, surprisingly, the prior observed advantage in the upper airway disappeared in recipient animals. The results suggest that furin site plays a role in efficiency of transmission and infection of recipient animals. Notably, despite clear attenuation relative to WT, the PQQAR mutant virus RNA was observed in every hamster pair, indicating transmission occurred even in the presence of WT SARS-CoV-2. Together, the results indicate that while the FCS is not strictly required, it does aid in transmission efficiency of SARS-CoV-2.

Overall, the manuscript data demonstrate that an intact furin cleavage site plays a critical role in SARS-CoV-2 infection, pathogenesis, and transmission efficiency. Disruption of the FCS in the S1/S2 loop results in ablated spike processing, altered host protease usage, and reduced disease in vivo. Importantly, while not strictly required for transmission, the FCS plays a critical role in efficiency of infection in recipient hamsters. Together, these results demonstrate the critical importance of an intact FCS in SARS-CoV-2 infection, pathogenesis, and transmission.

## Methods

### Cells

Vero E6 cells and Vero E6 cells expressing TMPRSS2 (Sekisui XenoTech) were grown in Dulbecco modified Eagle medium (DMEM; Gibco #11965–092) supplemented with 10% fetal bovine serum (FBS) (HyClone #SH30071.03) and 1% antibiotic-antimycotic (Gibco #5240062). Calu-3 2B4 cells were grown in DMEM supplemented with 10% FBS, 1% antibiotic-antimycotic, and 1 mg/mL sodium pyruvate. Human TMPRSS2 knockout Calu-3 cells (Abcam #273734) were grown in DMEM supplemented with 20% defined FBS (HyClone #SH30070.03), 1% antibiotic-antimycotic, 1% non-essential amino acid solution (NEAA; Gibco #11140050), and 1 mg/mL sodium pyruvate.

### Viruses

The recombinant WT and mutant SARS-CoV-2 virus sequences are based on the USA-WA1/2020 isolate sequence provided by the World Reference Center for Emerging Viruses and Arboviruses (WRCEVA), which was originally obtained from the US Centers for Disease Control and Prevention (CDC)^21^. Wild-type and mutant SARS-CoV-2 were generated using standard cloning techniques and reverse genetics system as previously described^22,23^ and propagated on Vero E6 cells. The mutations in PQQAR mutant have been verified by Sanger sequencing. The recovered mutant virus was further sequenced with NGS to confirm the maintenance of nucleotide mutations up to P2.

Infectious titers were measured by focus-forming assay. All experiments involving infectious virus were conducted at the University of Texas Medical Branch (UTMB) in an approved biosafety level (BSL) 3 laboratory with routine medical monitoring of staff.

### *In Vitro* Infection

Viral infections in Vero E6, TMRPSS2-expressing Vero E6, Calu-3 2B4, and Calu-3 TMPRSS2 KO cells were performed as previously described^22,24^. Briefly, cells were washed with phosphate buffered saline (PBS) and infected with WT or mutant SARS-CoV-2 at an MOI of 0.01 for 45 min at 37 °C. Following absorption, cells were washed three times with PBS and fresh growth media was added to represent time 0. Three or more biological replicates were collected at each time point and each experiment was repeated at least twice.

### Focus Forming Assay

Focus forming assays (FFAs) were performed as previously described^25^. Briefly, Vero E6 cells were seeded in 96-well plates to be 100% confluent. Samples were 10-fold serially diluted in serum-free media and 20 µl was to infect cells. Cells were incubated for 45 min at 37°C with 5% CO_2_ before 100 µl of 0.85% methylcellulose overlay was added. Cells were incubated for 24 hrs at 37°C with 5% CO_2_. After incubation, overlay was removed, and cells were washed three times with PBS before fixed and virus inactivated by 10% formalin for 30 min at room temperature. Cells were then permeabilized and blocked with 0.1% saponin/0.1% BSA in PBS before incubated with α-SARS-CoV-2 Nucleocapsid primary antibody (Cell Signaling Technology) at 1:1000 in permeabilization/blocking buffer overnight at 4°C. Cells are then washed three times with PBS before incubated with Alexa FluorTM 555-conjugated α-mouse secondary antibody (Invitrogen #A28180) at 1:2000 in permeabilization/blocking buffer for 1 h at room temperature. Cells were washed three times with PBS. Fluorescent foci images were captured using a Cytation 7 cell imaging multi-mode reader (BioTek), and foci were counted manually.

### Virion Purification and Western Blotting

Vero E6 cells were infected with WT or mutant SARS-CoV-2 at an MOI of 0.01. Supernatant was harvested 24 hpi and clarified by low-speed centrifugation. Virus particles were then pelleted by ultracentrifugation through a 20% sucrose cushion at 26,000 rpm for 3 h using a Beckman SW28 rotor. Pellets were resuspended in 2× Laemmli buffer to obtain protein lysates. Relative viral protein levels were determined by sodium dodecyl sulfate-polyacrylamide gel electrophoresis (SDS-PAGE) followed by Western blot analysis as previously described^11,12^. In brief, sucrose-purified WT and mutant SARS-CoV-2 virions were inactivated by boiling in Laemmeli buffer. Samples were loaded in equal volumes into 4 to 20% Mini-PROTEAN TGX Gels (Bio-Rad #4561093) and electrophoresed by SDS–PAGE. Protein was transferred to polyvinylidene difluoride (PVDF) membranes. Membranes were probed with SARS-CoV S-specific antibodies (Novus Biologicals #NB100-56578) and followed with horseradish peroxidase (HRP)-conjugated anti-rabbit antibody (Cell Signaling Technology #7074). Membranes were stripped and reprobed with SARS-CoV *N*-specific antibodies (Novus Biologicals #NB100-56576) and the HRP-conjugated anti-rabbit secondary IgG to measure loading. Signal developed using Clarity Western ECL substrate (Bio-Rad #1705060) or Clarity Max Western ECL substrate (Bio-Rad #1705062) and imaging on a ChemiDoc MP System (Bio-Rad #12003154). Densitometry was performed using ImageLab 6.0.1 (Bio-Rad #2012931).

### Hamster Infection Study

Male golden Syrian hamsters (3 to 4 weeks old) were purchased from Envigo (HsdHan:AURA strain). All studies were conducted under a protocol approved by the UTMB Institutional Animal Care and Use Committee and complied with USDA guidelines in a laboratory accredited by the Association for Assessment and Accreditation of Laboratory Animal Care. Procedures involving infectious SARS-CoV-2 were performed in the Galveston National Laboratory ABSL3 facility. Hamsters were intranasally inoculated with 10^5^ pfu of WT or PQQAR SARS-CoV-2 in100 µl. Infected hamsters were weighed and monitored for illness over 7 days. Hamsters were anesthetized with isoflurane (Henry Schein Animal Health) and nasal washes were collected with 400 µl of PBS on endpoint days (2, 4, and 7 dpi). Hamsters were euthanized by CO_2_ for organ collection. Nasal wash and lung were collected to measure viral titer. Left lungs were collected for histopathology.

### Hamster Transmission

Male golden Syrian hamsters (3-4 weeks old) were purchased from Envigo. Each group (WT, PQQAR, or PBS) had 10 donor hamsters that were intranasally infected with 100 uL of 10^5^ pfu of virus or PBS depending on the group. 24 hrs post infection, donor hamsters were cohoused 1:1 with a recipient hamster for 8 hrs for contact transmission. After 8 hrs, hamster pairs were separated into individual housing and the donors were nasal washed. At 2 days post infection for donors and post contact for recipients, hamsters were nasal washed with 400 µl of PBS and euthanized for lung and nasal wash collection. Nasal washes and lungs were processed in TRIzol, and RNA was extracted to perform next generation sequencing as previously described.

### Hamster Transmission Competition

Male golden Syrian hamsters (3-4 weeks old) were purchased from Envigo. ten donor hamsters were intranasally infected with a 1:1 ratio of WT:PQQAR SARS-CoV-2 totaling 10^5^ pfu in 100 µl and were subsequently individually housed. At 24 hrs post infection, donor hamsters were cohoused 1:1 with a recipient hamster for 12 hrs for contact transmission. After 8 hrs, hamster pairs were separated into individual housing and the donors were nasal washed for D1 values. At 2 and 4 days post infection for donors and post contact for recipients, hamsters were nasal washed with 400 µl of PBS and euthanized for lung and nasal wash collection. Nasal washes and lungs were processed in TRIzol. RNA was extracted to perform next generation sequencing.

### Histology

Histopathology was performed as previously described^26,27^. Briefly, left lungs were harvested from hamsters and fixed in 10% buffered formalin solution for at least 7 d. Fixed tissue was then embedded in paraffin, cut into 5 µM sections, and stained with hematoxylin and eosin on a SAKURA VIP6 processor by the University of Texas Medical Branch Surgical Pathology Laboratory. H&E staining was performed by the University of Texas Medical Branch Histology Laboratory and then analyzed and scored by a blinded pathologist.

### Immunohistochemistry

Antigen staining was performed as previously described^28^, Briefly, fixed and paraffin-embedded left lung lobes from hamsters were cut into 5 µM sections and mounted onto slides by the University of Texas Medical Branch Surgical Pathology Laboratory. Paraffin-embedded sections were warmed at 56°C for 10 min, deparaffinized with xylene (3x 5-min washes) and graded ethanol (3x 100% 5-min washes, 1x 95% 5-min wash), and rehydrated in distilled water. After rehydration, antigen retrieval was performed by steaming slides in antigen retrieval solution (10 mM sodium citrate, 0.05% Tween-20, pH 6) for 40 min (boil antigen retrieval solution in microwave, add slides to boiling solution, and incubate in steamer). After cooling and rinsing in distilled water, endogenous peroxidases were quenched by incubating slides in TBS with 0.3% H_2_O_2_ for 15 min followed by 2x 5-min washes in 0.05% TBST. Sections were blocked with 10% normal goat serum in BSA diluent (1% BSA in 0.05% TBST) for 30 min at room temperature. Sections were incubated with primary anti-N antibody (Sino #40143-R001) at 1:1000 in BSA diluent overnight at 4°C. Following overnight primary antibody incubation, sections were washed 3x for 5 min in TBST. Sections were incubated in secondary HRP-conjugated anti-rabbit antibody (Cell Signaling Technology #7074) at 1:200 in BSA diluent for 1 hour at room temperature. Following secondary antibody incubation, sections were washed 3x for 5 min in TBST. To visualize antigen, sections were incubated in ImmPACT NovaRED (Vector Laboratories #SK-4805) for 3 min at room temperature before rinsed with TBST to stop the reaction followed by 1x 5-min wash in distilled water. Sections were incubated in hematoxylin for 5 min at room temperature to counterstain before rinsing in water to stop the reaction. Sections were dehydrated by incubating in the previous xylene and graded ethanol baths in reverse order before mounted with coverslips.

### Structural Modeling

Structural models previously generated were used as a base to visualize residues mutated in Omicron^12^. Briefly, structural models were generated using SWISS-Model to generate homology models for WT and PQQAR SARS-CoV-2 spike protein based on the SARS-CoV-1 trimer structure (Protein Data Bank code 6ACD). Homology models were visualized and manipulated in PyMOL (version 2.5.4) to visualize the PQQAR mutation.

### Next Generation Sequencing and data analysis

Next generation sequencing (NGS) method was used to determine viral RNA populations from infected animals. Briefly, total cellular RNA samples were extracted from animal tissues and NGS libraries were prepared with Tiled-ClickSeq method^17,25^. A modified pre-RT annealing protocol was applied as previously described^25^ to reduce mis-priming. The final libraries comprising of 300–600 bps fragments were pooled and sequenced on an ElementBio Aviti platform with paired-end sequencing (120 bp R1 and 30 bp R2). The raw Illumina data of the Tiled-ClickSeq libraries were processed with established bioinformatics pipelines (https://github.com/andrewrouth/TCS). The relative ratio between WT and PQQAR was calculated based on the average Pilon-reported^29^ G-to-A mutation rate at specific genomic loci (G23607A, G23608A, G23610A, G23611A). A baseline mutation rate is also evaluated by averaging the background mutation rates from nts. 23606-23612.

## Acknowledgements

Special thanks to UTMB next generation sequencing core staff (Haiping Hao, Jill K. Thompson) for next-generation sequencing support.

## Author contributions

Conceptualization: ALM, MNV, VDM

Formal analysis: MNV, ALM, YZ, VDM

Funding acquisition: MNV, MSS, BAJ, KSP, VDM

Investigation: ALM, MNV, YZ, WMM, REA, KGL, YPA, LKE, JAP, DHW, KSP, VDM

Methodology: ALM, MNV, YZ, KGL, VDM

Project Administration: MNV, ALM, VDM

Supervision: MSS, DHW, KSP, VDM

Visualization: ALM, MNV, DHW, VDM

Writing – original draft: ALM, VDM

Writing – review and editing: ALM, MNV, KGL, DHW, VDM

